# DesignMaster: A Multi-Conditional Diffusion Framework for Rational PROTAC Design

**DOI:** 10.64898/2026.06.15.732318

**Authors:** Binze Shi, Jie Liu, Tong Pan, Yi Hao, Luke Isbel, Michael Roy, Ashley P. Ng, Xuequn Shang, Fuyi Li

## Abstract

**Motivation:** Proteolysis-targeting chimeras (PROTACs) enable targeted protein degradation through ternary complex formation with E3 ubiquitin ligase. However, the rational design of PROTACs remains highly challenging due to limited structure–activity relationship data and the vast conformational diversity of linkers. Existing computational approaches can be broadly divided into structure-based ternary modelling methods and fragment-based linker generation models. Although these approaches have advanced PROTAC design, they typically neglect key physicochemical constraints and linker-length control during the generation process, causing the generated PROTACs to lack balanced structural properties required for effective ternary complex formation with drug-like characteristics.

**Results:** To address these limitations, we propose DesignMaster, a diffusion-based generative framework that explicitly incorporates linker length and physicochemical properties as controllable conditioning signals. DesignMaster employs an E(3)-equivariant graph Transformer with a gated multi-condition fusion module to inject linker length and physicochemical constraints throughout the diffusion process, enabling fine-grained and constraint-aware molecular generation. Experiments on PROTAC-DB 2.0 and 3.0 demonstrate that DesignMaster outperforms state-of-the-art baselines, with a 3.2% improvement in validity and a 34.4% improvement in recovery. The Case study shows DesignMaster achieves a 51.78% reduction in RMSD when predicting the linker of PROTAC BCPyr targeting 6W7O, highlighting its potential for practical structure-guided PROTAC design.

**Availability:** The source code and datasets are available at https://github.com/ABILiLab/DesignMaster.

## Introduction

Proteolysis-targeting chimeras (PROTACs), first introduced by (Sakamoto et al., 2001), have emerged as a transformative modality for targeted protein degradation (Burslem and Crews, 2020; Zheng et al., 2022). A PROTAC is a heterobifunctional molecule composed of three modular components: a ligand that binds a protein of interest (POI), a linker, and a ligand that recruits an E3 ubiquitin ligase (Békés et al., 2022; Zhong et al., 2024; Li et al., 2022a). By simultaneously engaging the POI and the E3 ligase, PROTACs promote formation of a ternary POI–PROTAC–E3 complex, thereby hijacking the cellular ubiquitin–proteasome system to induce polyubiquitination and subsequent degradation of the target protein (Liu et al., 2025; Arnold, 2024). Distinct from conventional occupancy-driven inhibitors, PROTACs function through an event-driven and catalytic mechanism, enabling a single degrader molecule to participate in multiple rounds of target degradation at relatively low cellular concentrations. This makes PROTACs a highly promising strategy in anticancer drug development, particularly for targeting previously intractable cancer-associated proteins (Zhao and Dekker, 2022; Li et al., 2022b).

The rational design of PROTACs has remained inherently challenging since their initial inception, owing to their synthetic and mechanistic complexity and the limited availability of detailed structure–activity and experimental data, particularly in the early development of the field. (Zheng et al., 2022; Tran et al., 2022). Moreover, the degradation efficiency of a given PROTAC is critically governed by the geometry and dynamics of the ternary complex, which are strongly influenced by linker length, composition, and flexibility (Paiva and Crews, 2019). The vast conformational diversity arising from different linker designs, together with the many possibilities for POI/E3 binders and exit vectors, makes rational PROTAC optimisation a time-consuming and chemistry-intensive task, particularly in the absence of experimental structural data. Efficently exploring linker designs remains a central bottleneck in PROTAC development and motivates the need for systematic, structure-aware computational generation (Krieger et al., 2023).

In recent years, various computational methods have emerged to support rational PROTAC generation, in part building on a limited but growing literature of structural and biophysical datasets. These methods can be broadly categorised into two classes: structure-based ternary complex modelling approaches and fragment-based linker generation approaches (Li et al., 2024). The first class focuses on explicitly constructing and evaluating the POI–PROTAC–E3 ternary complex using molecular docking (Zheng et al., 2022; Nori et al., 2022), molecular dynamics (MD) simulations (Ma et al., 2025), or scoring function-based protocols (Bai et al., 2021; Weng et al., 2021). While such physics-driven approaches provide mechanistic interpretability and valuable structural insight, they are computationally intensive and often lack standardised protocols for reliable ternary modelling. These limitations have motivated the development of a second class of methods that recast PROTAC design as a fragment-connection problem. In this case, the warhead and E3 ligand are treated as predefined fragments, and the central task is to generate an appropriate linker to connect them. For example, DeLinker (Imrie et al., 2020) and 3DLinker (Huang et al., 2022) employ graph generative models to construct chemically valid linkers based on the spatial arrangement between two fragments. DiffLinker Igashov et al. (2024), DiffPROTACs Li et al. (2024), and DiffSBDD Schneuing et al. (2024) leverage diffusion-based 3D generative modelling to generate linkers that are geometrically consistent between fragment anchor positions.

Despite their notable performance, existing PROTAC generation models suffer from a major limitation: diffusion-based frameworks generally do not explicitly incorporate key physicochemical constraints or linker-length restrictions during the generation process. Consequently, the sampled PROTACs may be geometrically and chemically valid yet unconstrained in properties such as molecular weight, polarity, or flexibility. Balancing binding and geometry for productive ternary complex formation (Roy et al., 2019) with physicochemical profiles to enable more favourable drug-like characteristics, including aqueous solubility, cell permeability, and stability, is therefore critical (Hornberger and Araujo, 2023; Inganas et al., 2025). The absence of constraints around these properties can limit the practical applicability of current generative approaches.

To address this, we propose DesignMaster, a diffusion-based generative framework that explicitly incorporates linker length and physicochemical properties as controllable conditioning signals. DesignMaster first represents PROTAC molecules using an E(3)-equivariant graph Transformer to capture their three-dimensional structural features. We then design a gated multi-condition fusion module to inject conditional information into every stage of the diffusion denoising trajectory, enabling fine-grained control over the generative process. Utilizing PROTAC-DB demonstrates state-of-the-art performance in terms of validity and recovery (See Supplementary section S2 for the definitions of the metrics). Furthermore, DesignMaster achieves a 51.78% decrease in RMSD between the generated PROTAC and the ground-truth PROTAC BCPyr (PDB: 6W7O), demonstrating its effectiveness for practical structure-guided PROTAC development. These results highlight the potential of DesignMaster as a practical computational tool to support controllable and property-aware PROTAC design.

## Materials and Methods

### Data Collection and Preprocessing

Following (Li et al., 2024), we conduct experiments on three complementary datasets: GEOM (Axelrod and Gomez-Bombarelli, 2022), PROTAC-DB 2.0 (Weng et al., 2023), and PROTAC-DB 3.0 (Ge et al., 2025). GEOM provides large-scale 3D molecular conformations and is used to pretrain the E(3)-equivariant GNN backbone, enabling robust geometric representation learning prior to fine-tuning on relatively small PROTAC datasets.

For downstream evaluation, we use PROTAC-DB 2.0 and the expanded PROTAC-DB 3.0 (Weng et al., 2023; Ge et al., 2025). Since PROTAC entries are provided in 2D format, we computationally generate 3D conformations and decompose each molecule into warhead, E3 ligand, and linker via graph-based substructure matching. Samples that cannot be reliably decomposed are removed.

Following prior work (Li et al., 2024), we adopt the same preprocessing and data-splitting protocol for both PROTAC-DB versions to ensure a fair comparison. Each dataset is independently partitioned into training, validation, and test sets with a 5:1:1 ratio. Full preprocessing details are provided in Supplementary Section S1.

### The Framework of DesignMaster

An overview of the architecture of DesignMaster, and its key components are visualised in Fig. 1. DesignMaster is a diffusion-based generative model that explicitly incorporates linker length and physiochemical properties as controllable conditioning signals. It consists of four major components: diffusion network, denoising network, equivariant graph nerual network, and gated multi-condition fusion. These four components will be elaborated in the following sections.

**Fig. 1.**
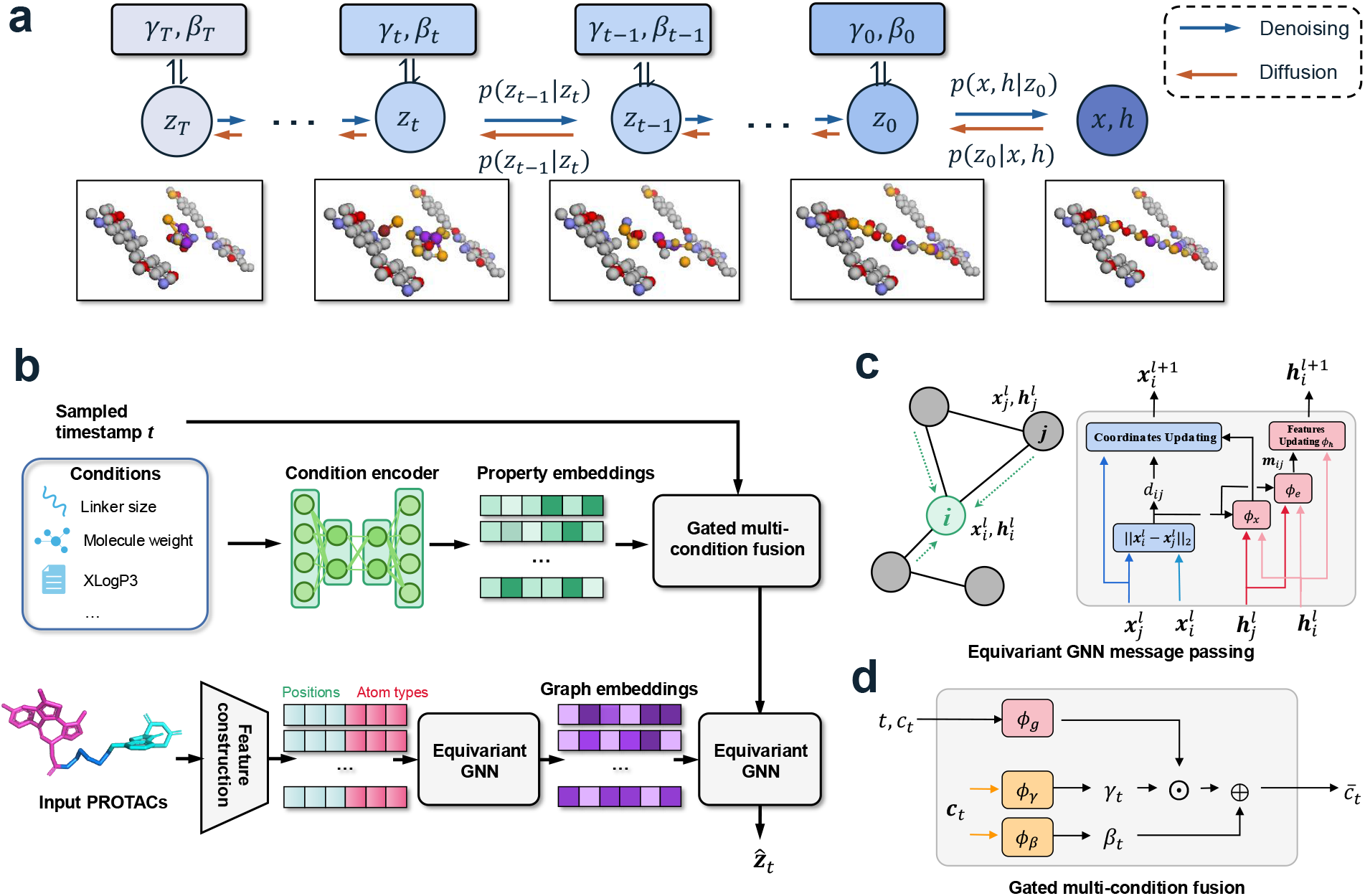
Overview of the DesignMaster framework. **a**. Forward diffusion and reverse denoising process for conditional PROTAC generation, in which the warhead and E3 ligand remain fixed in both structure and spatial position, while the noisy latent variables **z**_*t*_ are progressively denoised to synthesize the linker, resulting in the final molecule structure (**x, h**). **b**. Overall model architecture. Molecular graphs and multi-property conditions (e.g., linker size, molecular weight, XLogP3) are encoded and fused via a gated multi-condition module to guide the equivariant generative process. **c**. E(3)-equivariant GNN message passing, where atomic coordinates and features are jointly updated to preserve geometric equivariance. **d**. Gated multi-condition fusion mechanism that modulates the diffusion dynamics by integrating conditional signals into the latent variables at each timestep.

#### Diffusion Process

We adopt a denoising diffusion probabilistic model (DDPM) to generate molecular structures in 3D space, following the formulation introduced by (Hoogeboom et al., 2022). As depicted in Fig. 1a, the model operates on a latent variable **z**_*t*_, which represents the noisy molecular configuration at diffusion timestep *t*. Each **z**_*t*_ consists of atomic coordinates **x**_*t*_ ∈ ℝ^*N* ×3^ and associated node features **h**_*t*_ ∈ ℝ^*N* ×*nf*^, where *N* is the number of atoms and *nf* is the dimension of node features.

The forward diffusion process gradually perturbs the clean molecular representation **z**_0_ by injecting Gaussian noise over *T* timesteps according to a predefined noise schedule 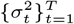:

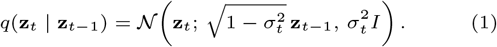

This process admits a closed-form marginal distribution:

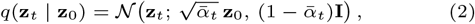

where 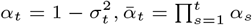.

The reverse diffusion process aims to progressively denoise **z**_*t*_ and recover a valid molecular structure conditioned on molecular context and external constraints. It is defined as a parameterized Gaussian transition:

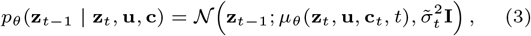

where 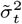 denotes the posterior variance. the mean is parameterized via a noise prediction network *ϵ*_*θ*_:

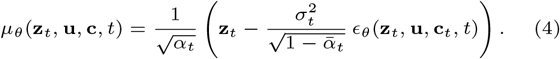

Here, **u** denotes a mask matrix preventing fixed warhead and E3 ligand being disturbed, while **c**_*t*_ represents additional condition encoding information used to guide the diffusion process.

The denoising network is trained to predict the Gaussian noise added at each diffusion step. Specifically, a timestep *t* ∼ *U*{(1, …, *T*)} and noise *ϵ* ∼ *N*(**0, I**) are sampled, and the noisy input is constructed as:

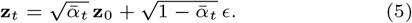

The training objective is then defined as:

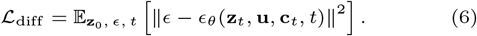

#### Denoising Network

We now describe the architecture used to parameterize the noise prediction function *ϵ*_*θ*_(**z**_*t*_, **u, c**, *t*). At diffusion timestep *t*, the noisy latent variable **z**_*t*_ is decomposed into atomic coordinates and node features **z**_*t*_ = (**x**_*t*_, **h**_*t*_). To extract geometry-aware representations, the pair (**x**_*t*_, **h**_*t*_) is first processed by an E(3)-equivariant graph neural network pretrained on the large-scale GEOM dataset:

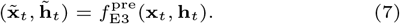

This pretrained encoder serves as a geometry feature extractor, providing stable and physically meaningful representations of noisy molecular conformations. In parallel, the external condition encoding **c** is modulated by a time-dependent gated fusion mechanism to produce a timestep-aware conditioning signal:

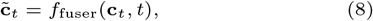

where *f*_fuser_ denotes the time-gated condition fuser. The fused condition features are then concatenated to the equivariant node features 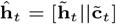, while the coordinate representation 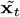 is preserved to maintain equivariance. The augmented representations 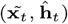 are subsequently passed to a second E(3)-equivariant GNN, which is trained from scratch and dedicated to denoising:

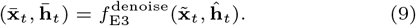

Rather than directly predicting noise, the model outputs a denoised estimate of the molecular representation. The predicted noise is then obtained by computing the residual between the input and output representations:

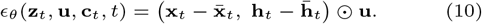

This residual formulation aligns with standard DDPM training objectives and allows the model to focus on learning the denoising displacement while preserving E(3)-equivariance.

#### Equivariant Graph Neural Network

We adopt an E(3)-equivariant graph neural network to encode and denoise molecular structures in 3D space (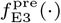 and 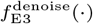). Let **x**_*t*_ ∈ ℝ^*N* ×3^ and **h**_*t*_ ∈ ℝ^*N* ×*nf*^ denote the coordinates and node features of atom *i*, respectively. Here, *nf* corresponds to the number of atom or molecular types, represented as a one-hot encoding. These features are subsequently projected into a higher-dimensional latent space within the network.

At layer *ℓ* pairwise messages are computed based on node features and relative geometry:

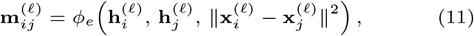

where *ϕ*_*e*_(·) is a fully connected neural network shared across all edges. The coordinate update follows the standard equivariant formulation:

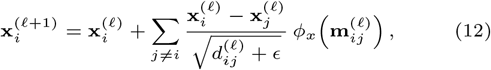

where 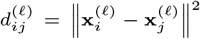, which ensures E(3)-equivariance by construction. The node features are updated by aggregating incoming messages:

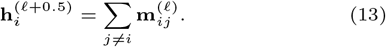

To enhance the expressiveness of node feature updates, we implement *ϕ*_*h*_(·) using a Graphormer-style attention mechanism rather than a simple MLP. Given the aggregated message 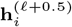, the node features are updated via self-attention over all nodes:

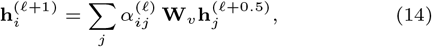

where **W**_*v*_ is a learnable linear projection and 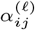 denotes the attention weight between nodes *i* and *j*. The attention weights are computed as:

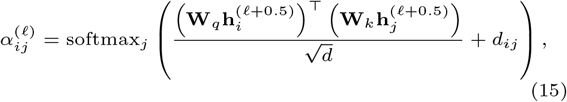

where **W**_*q*_ and **W**_*k*_ are learnable projection matrices, *d* denotes the hidden feature dimension.

#### Gated Multi-Condition Fusion

To incorporate conditional information into the diffusion process, we introduce a gated multi-condition fusion module *f*_fuser_, illustrated in Fig. 1d. This module is inspired by the Feature-wise Linear Modulation (FiLM) paradigm ((Perez et al., 2018)), where external conditions modulate the latent representation through learnable scaling and shifting parameters, while the diffusion timestep controls the strength of conditioning via a gating mechanism.

We first compute a timestep-dependent gating scalar (or vector) from the latent representation **g**_*t*_ = *ϕ*_*g*_(*t*), where *ϕ*_*g*_(·)is a learnable function that produces a gate controlling the overall influence of conditioning at timestep *t*. The condition encoding *c*_*t*_ is mapped to FiLM parameters via two learnable projections:

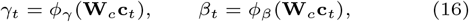

where *γ*_*t*_ and *β*_*t*_ denote feature-wise scaling and shifting coefficients, respectively.

The final fused representation is obtained by applying FiLM modulation to, gated by the timestep-dependent factor:

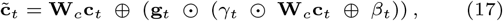

where ⊙ denotes element-wise multiplication and ⊗ denotes element-wise addition.

## Results and Discussion

We comprehensively evaluated the proposed framework from multiple perspectives, including model performance, property alignment, and distributional consistency.

### Model Performance

Table 1 reports the quantitative results on PROTAC-DB 2.0 and 3.0. Overall, our model achieves the best or highly competitive performance across all metrics. On PROTAC-DB 2.0, it outperforms all baselines in Validity (98.62%), Uniqueness (71.30%), and Recovery (57.50%). On PROTAC-DB 3.0, our method again achieves the highest Validity (97.01%) and Recovery (60.38%). Compared to the most competitive baseline DiffPROTACs, DesignMaster obtains a 3.2% performance gap on validity and 34.4% on recovery, demonstrating strong generalization to the newer and more diverse dataset.

**Table 1.** Performance comparison on PROTAC-DB 2.0 (V 2.0) and 3.0 (V 3.0). Validity: the proportion of chemically valid molecules generated by the model. Uniqueness: the fraction of non-duplicate molecules among the valid generated samples. Recovery: the percentage of reference molecules that are successfully reconstructed by the model.

| Dataset | Model | Validity(%) | Uniqueness(%) | Recovery(%) |
| --- | --- | --- | --- | --- |
| V2.0 | EDM | 91.35 | 48.54 | 35.25 |
|  | DiffLinker | 92.10 | 56.80 | 39.20 |
|  | DiffPROTACs | 93.86 | 68.70 | 43.75 |
|  | DesignMaster | **98.62** | **71.30** | **57.50** |
| V3.0 | EDM | 91.72 | 45.10 | 33.80 |
|  | DiffLinker | 92.60 | 54.20 | 40.75 |
|  | DiffPROTACs | 93.96 | **67.74** | 44.91 |
|  | DesignMaster | **97.01** | 60.49 | **60.38** |
|  | DesignMaster (w/o CE) | 94.93 | 57.01 | 59.62 |
|  | DesignMaster (w/o GF) | 95.58 | 54.72 | 62.29 |

Compared with the baselines, the performance gains can be attributed to both architectural and task-specific design choices. EDM relies on an MLP-based message-passing scheme, which is less expressive than our Graphormer-enhanced equivariant network for capturing complex structural dependencies. DiffLinker is designed as a general-purpose linker generator and does not explicitly model the unique multi-component structure of PROTACs. Although DiffPROTACs adapts diffusion to the PROTAC setting, it does not explicitly encode rich molecular attributes during denoising. In contrast, our framework incorporates conditional molecular feature encoding, enabling more chemically informed generation. The consistently higher Recovery, especially on PROTAC-DB 3.0, highlights the importance of explicitly modeling multi-level molecular properties for faithful linker reconstruction.

The ablation results further confirm the effectiveness of each component. Removing the conditional encoding (w/o CE) degrades overall performance, indicating the necessity of molecular attribute guidance. Similarly, removing the time-gated fusion mechanism (w/o GF) leads to performance fluctuations, showing that properly balancing structural and conditional information during denoising is critical for stable and accurate generation.

### Property Alignment

We evaluate the alignment between generated and real PROTAC molecules by comparing the distributions of key physicochemical descriptors, including Hydrogen Bond Acceptor Count (HBA), Hydrogen Bond Donor Count (HBD), rotatable bond count (RB), XLogP3, heavy atom count (HA), and Topological Polar Surface Area (TPSA). As shown in Fig. 2, the generated molecules closely follow the empirical distributions of the reference datasets, with substantial overlap observed for size-related (molecular weight, HA) and polarity-related (TPSA). Minor deviations appear in distribution tails, but overall distributional shapes are well preserved. The complete property distributions are provided in Supplementary Fig. S1.

**Fig. 2.**
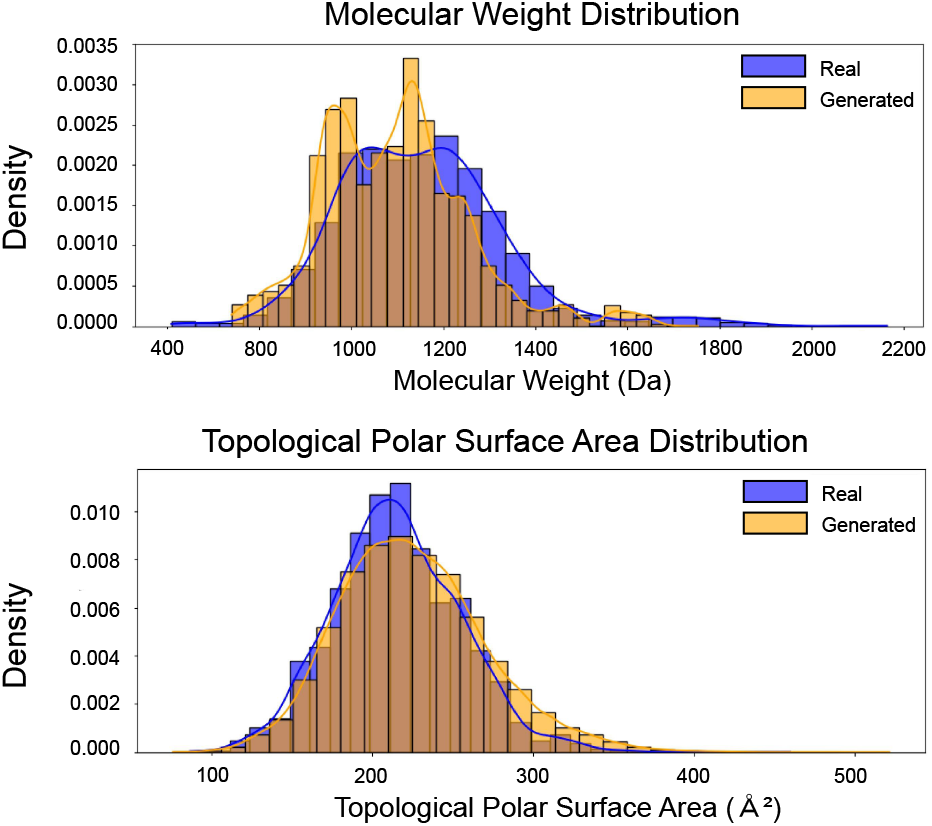
Property distribution gap between generated and true linkers.

To quantify the impact of multi-condition encoding, we compute the Wasserstein distances between the original and reconstructed property distributions, as shown in Fig. 3 and Table 2. The comparison between the full model and its ablated variant clearly highlights the contribution of conditional modulation.

**Table 2.** Distances between original and reconstructed property distributions. Values are reported as mean ± standard deviation over five independent runs. w/o mc indicates without multi-condition module.

| Model | HBA | HBD | RB | XLogP3 | HA | TPSA |
| --- | --- | --- | --- | --- | --- | --- |
| w/o mc (3.0) | 3.68 ± 0.15 | 4.25 ± 0.10 | 6.08 ± 0.20 | 4.05 ± 0.18 | 0.55 ± 0.05 | 9.32 ± 0.35 |
| Full (2.0) | 2.18 ± 0.12 | 4.29 ± 0.08 | 0.45 ± 0.07 | 4.34 ± 0.15 | 0.57 ± 0.04 | 5.28 ± 0.25 |
| Full (3.0) | 2.36 ± 0.14 | 4.26 ± 0.09 | 0.83 ± 0.09 | 4.29 ± 0.14 | 0.52 ± 0.04 | 3.66 ± 0.22 |

**Fig. 3.**
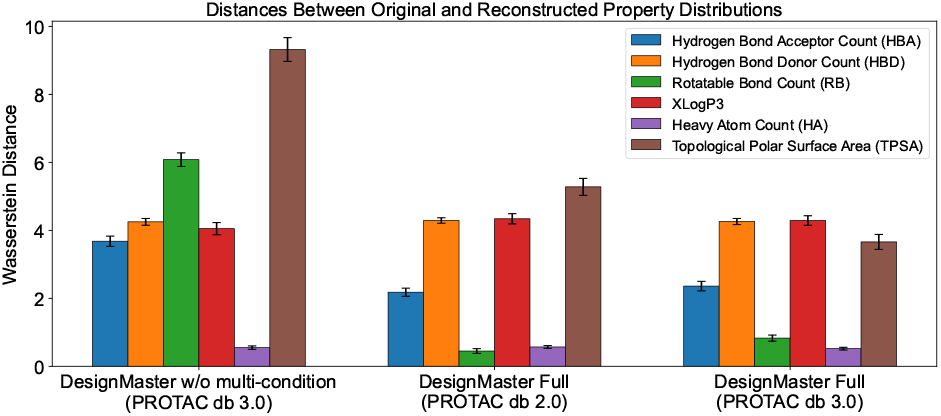
Comparison of Wasserstein distances across DesignMaster variants. Results are reported as the mean over five independent runs, with standard deviations indicated as error bars.

When the multi-condition module is removed (DesignMaster w/o multi-condition, trained on PROTAC-DB 3.0), the distribution gaps increase substantially for several key properties, particularly HBA (3.68), RB (6.08), and TPSA (9.32). These properties are closely related to hydrogen-bonding capacity, conformational flexibility, and polarity, suggesting that without explicit conditional guidance, the model has greater difficulty preserving the physicochemical characteristics of realistic PROTAC linkers.

In contrast, the full model trained on PROTAC-DB 2.0 significantly reduces the gaps for HBA (2.18), RB (0.45), and TPSA (5.28), indicating that multi-condition modulation effectively constrains flexibility and polar surface properties during generation. The trend remains consistent when trained on PROTAC-DB 3.0, where the TPSA gap further decreases to 3.66 and RB remains relatively low (0.83), demonstrating stable generalization under a larger and more diverse dataset.

For heavy atom count (HA), all variants achieve comparable results, as expected since molecular size is a fundamental structural attribute that is typically well constrained or implicitly regularized in existing generative frameworks. The hydrogen bond donor count (HBD) also remains nearly identical across models, which is reasonable given that this attribute is often carefully controlled in PROTAC design due to its impact on physicochemical properties such as cell permeability (Hornberger and Araujo, 2023). XLogP3 shows only minor differences among variants.

Overall, both the distribution plots and the Wasserstein distance values demonstrate that our property-aware diffusion framework generates linkers whose physicochemical distributions are better aligned with the true PROTAC linker space, highlighting the effectiveness of explicit molecular attribute encoding.

## Case Study

### Evaluation of Linker Reconstruction

To assess the structural validity of generated linkers, we fix the warhead and E3 ligand fragments and generate multiple linker candidates using DesignMaster and DiffPROTACs, respectively. The assembled PROTAC molecules are then subjected to molecular docking using AutoDock Vina against the target ternary complex structure (PDB: 6W7O). The resulting docking poses are compared to the experimentally resolved PROTAC conformation by calculating the RMSD.

As shown in Fig. 4, DesignMaster consistently produces linker-generated PROTACs that recover near-native binding geometries, achieving RMSD values of 0.43, 0.56, and 0.64 for the top-3 ranked docking poses. In contrast, DiffPROTACs exhibits larger deviations (0.65, 1.19, and 1.54 Å), with substantial pose drift observed in lower-ranked solutions. Notably, DesignMaster demonstrates both lower top-1 RMSD and improved top-k stability, indicating stronger structural coherence across generated linkers.

**Fig. 4.**
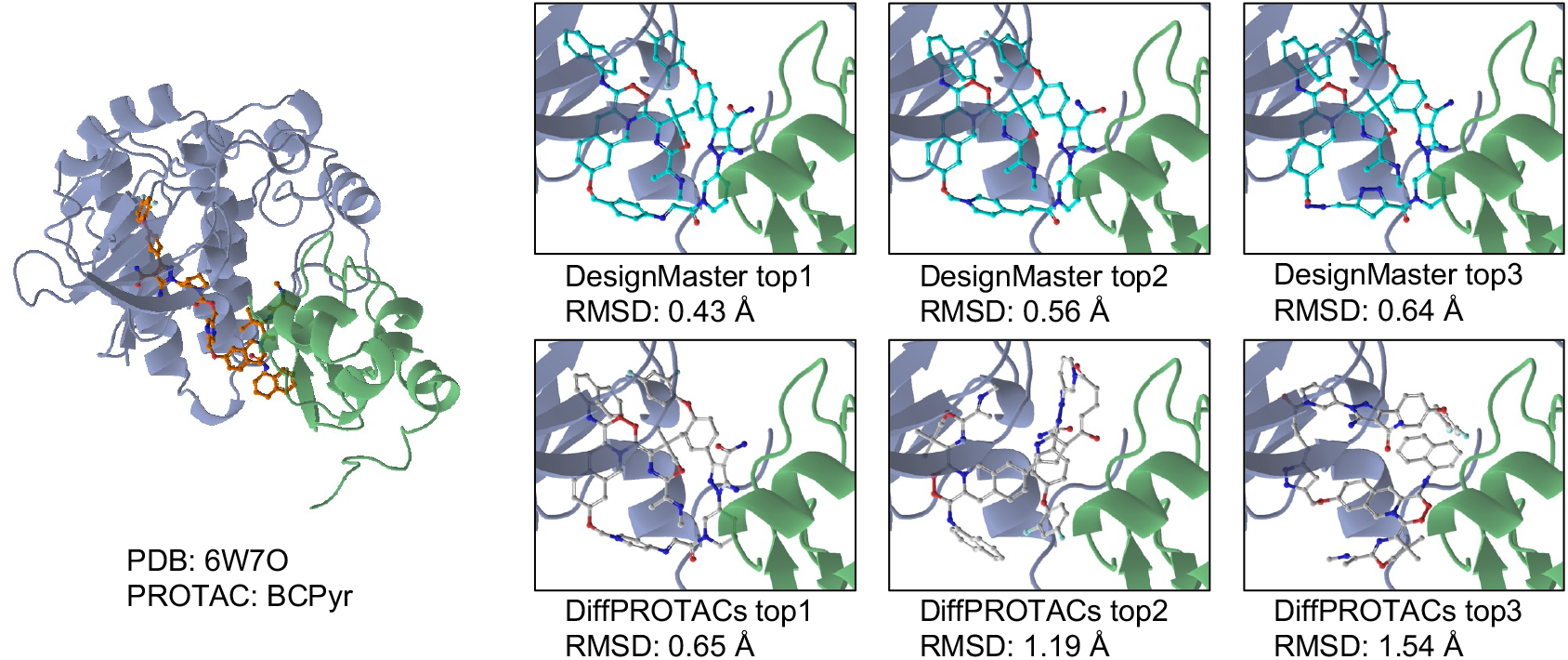
Case study. Evaluation of linker generation on PDB 6W7O (PROTAC: **BCPyr**). Linkers are generated by DesignMaster and DiffPROTACs, and RMSD values are computed between the generated PROTAC and the reference PROTAC conformation.

Qualitative observations from Fig. 4 suggest that DesignMaster better preserves the spatial orientation between the warhead and E3 ligand, resulting in closer alignment with the reference conformation. DiffPROTACs, by comparison, shows increased conformational distortion, particularly in distal linker regions, leading to suboptimal binding alignment.

These results suggest that the geometry-aware equivariant backbone and multi-condition modulation mechanism in DesignMaster enhance the structural realism of generated linkers, translating into improved recovery of native ternary binding conformations.

## Denoising Dynamics Analysis

Fig. 5 compares the denoising trajectories of DiffPROTACs and our model by tracking total loss, position loss, and feature loss across sampling steps. Both methods exhibit a general decreasing trend as the diffusion process progresses, consistent with iterative noise removal. However, our model demonstrates a more stable and consistently downward trajectory, particularly in the later sampling stages. The total loss decreases more smoothly and reaches a lower final magnitude, indicating improved convergence behavior. In contrast, DiffPROTACs shows more pronounced oscillations and a slower decline in the later timesteps, suggesting less stable denoising dynamics.

**Fig. 5.**
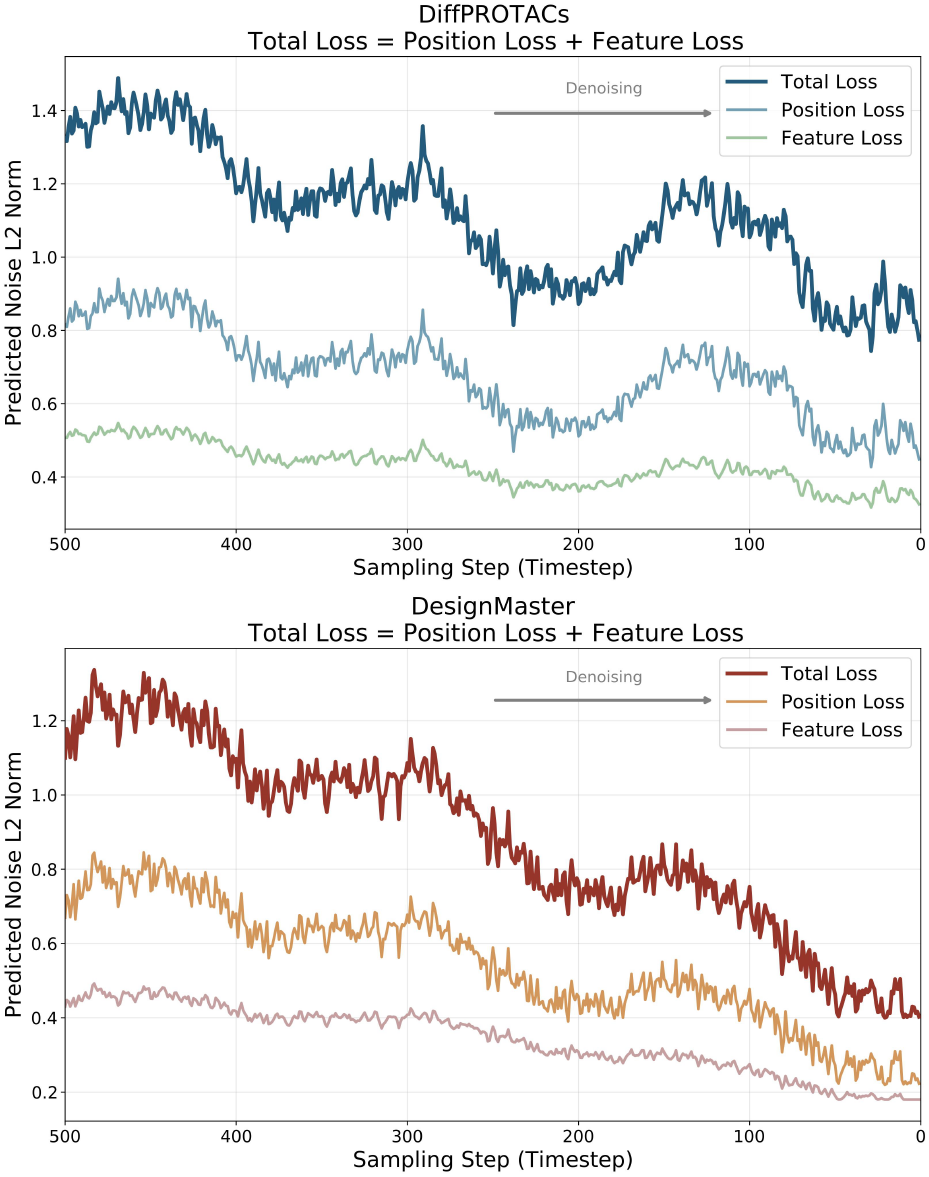
Comparison of denoising trajectories between DiffPROTACs and DesignMaster, showing total, position, and feature loss trends across time steps.

Notably, the position and feature losses in our model decrease in a coordinated manner, implying that geometric reconstruction and chemical feature optimization progress in a balanced way. This coordinated denoising behavior translates into more stable structural refinement, reducing abrupt conformational shifts during generation. For biological applications, such stability is critical, as it improves reproducibility and reduces the likelihood of generating structurally inconsistent or physically unrealistic linker conformations.

One possible explanation for this improved stability is the incorporation of linker-length and physicochemical constraints, which effectively restrict the sampling space and guide the model toward chemically plausible regions. While these constraints reduce sampling freedom, they act as a beneficial regularization mechanism, preventing excessive structural drift and promoting more reliable convergence.

## Conclusion

In this work, we propose DesignMaster, a diffusion-based framework for constraint-aware PROTAC linker design. By incorporating linker length and physicochemical properties as conditioning signals within an E(3)-equivariant graph Transformer, the model enables geometry-aware and controllable molecular generation.

Experiments on PROTAC-DB 2.0 and 3.0 show that DesignMaster outperforms existing methods, achieving improved validity and recovery. Docking-based evaluation further demonstrates enhanced structural accuracy, with a 51.78% reduction in RMSD in a case study on IAP-recruiting Bruton’s Tyrosine Kinase (BTK) PROTAC **BCPyr** (PDB: 6W7O). These results indicate the potential of conditional diffusion modeling for structure-guided PROTAC design.

Building on this controllable generative framework, our future studies aim at adaptively reweighting linker property constraints based on the intrinsic physicochemical features of fixed POI and E3 binders, allowing targeted optimization of polarity, hydrogen bonding, and molecular size in a context-dependent manner.

## References

Arnold, C. (2024). Protac protein degraders to drug the undruggable enter phase 3 trials. Nat. Med, 30(11):3030–3031.

Axelrod, S. and Gomez-Bombarelli, R. (2022). Geom, energy-annotated molecular conformations for property prediction and molecular generation. Scientific Data, 9(1):185.

Bai, N., Miller, S. A., Andrianov, G. V., Yates, M., Kirubakaran, P., and Karanicolas, J. (2021). Rationalizing protac-mediated ternary complex formation using rosetta. Journal of chemical information and modeling, 61(3):1368–1382.

Békés, M., Langley, D. R., and Crews, C. M. (2022). Protac targeted protein degraders: the past is prologue. Nature reviews Drug discovery, 21(3):181–200.

Burslem, G. M. and Crews, C. M. (2020). Proteolysis-targeting chimeras as therapeutics and tools for biological discovery. Cell, 181(1):102–114.

Ge, J., Li, S., Weng, G., Wang, H., Fang, M., Sun, H., Deng, Y., Hsieh, C.-Y., Li, D., and Hou, T. (2025). Protac-db 3.0: an updated database of protacs with extended pharmacokinetic parameters. Nucleic acids research, 53(D1):D1510–D1515.

Hoogeboom, E., Satorras, V. G., Vignac, C., and Welling, M. (2022). Equivariant diffusion for molecule generation in 3d. In International conference on machine learning, pages 8867–8887. PMLR.

Hornberger, K. R. and Araujo, E. M. (2023). Physicochemical property determinants of oral absorption for protac protein degraders. Journal of Medicinal Chemistry, 66(12):8281–8287.

Huang, Y., Peng, X., Ma, J., and Zhang, M. (2022). 3dlinker: an e (3) equivariant variational autoencoder for molecular linker design. arXiv preprint arXiv:2205.07309.

Igashov, I., Stärk, H., Vignac, C., Schneuing, A., Satorras, V. G., Frossard, P., Welling, M., Bronstein, M., and Correia, B. (2024). Equivariant 3d-conditional diffusion model for molecular linker design. Nature Machine Intelligence, 6(4):417–427.

Imrie, F., Bradley, A. R., van der Schaar, M., and Deane, C. M. (2020). Deep generative models for 3d linker design. Journal of chemical information and modeling, 60(4):1983–1995.

Inganas, E., Lavén, H., Caraballo, R., Klingegard, F., Eklund, M., Andersson, L., Hilgendorf, C., and Matsson, P. (2025). Conformational dynamics in the cell membrane interactions of bispecific targeted degrader therapeutics. Journal of Medicinal Chemistry, 68(24):25881–25898.

Krieger, J., Sorrell, F. J., Wegener, A. A., Leuthner, B., Machrouhi-Porcher, F., Hecht, M., Leibrock, E. M., Müller, J. E., Eisert, J., Hartung, I. V., et al. (2023). Systematic potency and property assessment of vhl ligands and implications on protac design. ChemMedChem, 18(8):e202200615.

Li, F., Hu, Q., Zhang, X., Sun, R., Liu, Z., Wu, S., Tian, S., Ma, X., Dai, Z., Yang, X., et al. (2022a). Deepprotacs is a deep learning-based targeted degradation predictor for protacs. Nature communications, 13(1):7133.

Li, F., Hu, Q., Zhou, Y., Yang, H., and Bai, F. (2024). Diffprotacs is a deep learning-based generator for proteolysis targeting chimeras. Briefings in bioinformatics, 25(5).

Li, X., Pu, W., Zheng, Q., Ai, M., Chen, S., and Peng, Y. (2022b). Proteolysis-targeting chimeras (protacs) in cancer therapy. Molecular cancer, 21(1):99.

Liu, J., Roy, M. J., Isbel, L., and Li, F. (2025). Accurate protac-targeted degradation prediction with degrademaster. Bioinformatics, 41(Supplement 1):i342–i351.

Ma, N., Bhattacharya, S., Muk, S., Jandova, Z., Schmalhorst, P. S., Ghosh, S., Le, K., Diers, E., Trainor, N., Farnaby, W., et al. (2025). Frustration in the protein-protein interface plays a central role in the cooperativity of protac ternary complexes. Nature Communications, 16(1):8595.

Nori, D., Coley, C. W., and Mercado, R. (2022). De novo protac design using graph-based deep generative models. arXiv preprint arXiv:2211.02660.

Paiva, S.-L. and Crews, C. M. (2019). Targeted protein degradation: elements of protac design. Current opinion in chemical biology, 50:111–119.

Perez, E., Strub, F., De Vries, H., Dumoulin, V., and Courville, A. (2018). Film: Visual reasoning with a general conditioning layer. In Proceedings of the AAAI conference on artificial intelligence, volume 32.

Roy, M. J., Winkler, S., Hughes, S. J., Whitworth, C., Galant, M., Farnaby, W., Rumpel, K., and Ciulli, A. (2019). Spr-measured dissociation kinetics of protac ternary complexes influence target degradation rate. ACS chemical biology, 14(3):361–368.

Sakamoto, K. M., Kim, K. B., Kumagai, A., Mercurio, F., Crews, C. M., and Deshaies, R. J. (2001). Protacs: Chimeric molecules that target proteins to the skp1–cullin–f box complex for ubiquitination and degradation. Proceedings of the National Academy of Sciences, 98(15):8554–8559.

Schneuing, A., Harris, C., Du, Y., Didi, K., Jamasb, A., Igashov, I., Du, W., Gomes, C., Blundell, T. L., Lio, P., et al. (2024). Structure-based drug design with equivariant diffusion models. Nature Computational Science, 4(12):899–909.

Tran, N. L., Leconte, G. A., and Ferguson, F. M. (2022). Targeted protein degradation: design considerations for protac development. Current protocols, 2(12):e611.

Weng, G., Cai, X., Cao, D., Du, H., Shen, C., Deng, Y., He, Q., Yang, B., Li, D., and Hou, T. (2023). Protac-db 2.0: an updated database of protacs. Nucleic acids research, 51(D1):D1367–D1372.

Weng, G., Li, D., Kang, Y., and Hou, T. (2021). Integrative modeling of protac-mediated ternary complexes. Journal of Medicinal Chemistry, 64(21):16271–16281.

Zhao, C. and Dekker, F. J. (2022). Novel design strategies to enhance the efficiency of proteolysis targeting chimeras. ACS Pharmacology & Translational Science, 5(9):710–723.

Zheng, S., Tan, Y., Wang, Z., Li, C., Zhang, Z., Sang, X., Chen, H., and Yang, Y. (2022). Accelerated rational protac design via deep learning and molecular simulations. Nature Machine Intelligence, 4(9):739–748.

Zhong, G., Chang, X., Xie, W., and Zhou, X. (2024). Targeted protein degradation: advances in drug discovery and clinical practice. Signal transduction and targeted therapy, 9(1):308.

