## Supplementary Material for "DesignMaster: A Multi-Conditional Diffusion Framework for Rational PROTAC Design"

#### 1 Data Preprocessing

We conducted our experiments using three complementary datasets: GEOM, PROTAC-DB 2.0, and PROTAC-DB 3.0.

The GEOM dataset is a large collection of 3D molecular conformations generated via extensive sampling and energy annotation for diverse molecules. It contains hundreds of thousands of molecules with ensembles of low-energy conformers and associated structural information, and has been widely used as a benchmark for property prediction and generative modeling in 3D molecular space. In this work, we use the GEOM dataset to pretrain the E(3)-equivariant graph neural network that serves as the backbone feature extractor in our diffusion model. Pretraining on GEOM allows the network to learn meaningful geometric representations of noisy molecular structures, which is critical given the limited size of available PROTAC datasets.

PROTAC-DB 2.0 is a curated database of PROTAC molecules and their associated components, published as part of an effort to support data-driven research in targeted protein degradation. It contains structural and experimental information for 3,270 PROTACs, including 365 warheads, 82 E3 ligands, and 1,501 linkers. The database also includes multiple types of bioactivity and binding affinity data and a number of ternary complex structures, making it useful for both analysis and model evaluation. In our experiments, PROTAC-DB 2.0 serves as a baseline dataset for evaluating the performance of linker generation and reconstruction.

[†]These authors contributed equally to this work.

PROTAC-DB 3.0 is the most recent and comprehensive release of the PROTAC-DB database, significantly expanding on version 2.0. In this update, the number of PROTAC entries increases to 6,111, representing an approximately 87% growth over PROTAC-DB 2.0, with 569 warheads, 107 E3 ligands, and 2,753 linkers included. Importantly, PROTAC-DB 3.0 also integrates additional biological and pharmacokinetic annotations, as well as a larger set of ternary complex structures for protein–PROTAC–E3 interactions, which improves the diversity and utility of data for computational modeling.

Following prior work, we adopt the same preprocessing and data splitting protocol as for the PROTAC datasets. Since the PROTAC entries in PROTAC-DB are provided only in 2D format and lack experimentally resolved 3D structures, we computationally generated 3D conformations for all PROTAC molecules. Specifically, 3D structures were constructed using a random conformer generation strategy, which offers a practical trade-off between computational efficiency and structural diversity.

To decompose each PROTAC molecule into its constituent warhead, E3 ligand, and linker, we leverage the structural annotations provided by PROTAC-DB and perform subgraph matching between the full PROTAC and its components. This process is formulated as a subgraph isomorphism problem and solved using graph-based matching procedures, yielding explicit atom-level mappings between linkers and the full molecules. Samples that could not be reliably decomposed or lacked valid linker structures were discarded.

Since our experiments involve both PROTAC-DB 2.0 and PROTAC-DB 3.0, we follow the same preprocessing and splitting strategy for each version independently to ensure fair and consistent evaluation. For PROTAC-DB 2.0, we adopt the standard split used in DiffPROTAC. After preprocessing and filtering invalid samples, 2,813 PROTAC molecules remain and are randomly divided into training, validation, and test sets with a ratio of 5:1:1. For PROTAC-DB 3.0, which contains a significantly larger and more diverse collection of PROTAC molecules, we apply the same preprocessing pipeline and splitting protocol. The resulting dataset is independently partitioned into training, validation, and test sets using the same ratio as PROTAC-DB 2.0. This ensures that performance comparisons across the two datasets reflect differences in data scale and diversity rather than discrepancies in data processing or evaluation methodology.

#### 2 Metrics

To evaluate the quality of generated linkers and reconstructed PROTAC molecules, we adopt three standard metrics following DiffPROTAC: Validity, Uniqueness, and Recovery. Let $\mathcal{G} = \{\hat{L}_i\}_{i=1}^N$ denote the set of generated linkers, and $\mathcal{T} = \{L_i\}_{i=1}^N$ denote the corresponding ground-truth linkers in the test set.

**Validity** measures the proportion of chemically valid molecules among the generated samples. A generated linker is considered valid if it satisfies chemical valency rules and can be successfully parsed into a valid molecular graph with the help of OpenBabel and RDKit. Formally, validity is defined as:

$$\text{Validity} = \frac{1}{N} \sum_{i=1}^{N} \mathbf{1}\big(\hat{L}_i \text{ is chemically valid}\big), \tag{1}$$

**Recovery** measures the proportion of test samples for which the ground-truth linker is successfully reconstructed by the model. Specifically, for each test instance $i$, multiple linker candidates are generated, and recovery is defined as:

$$\text{Recovery} = \frac{1}{N} \sum_{i=1}^{N} \mathbf{1}\big(L_i \in \{\hat{L}_i^{(k)}\}_{k=1}^K\big), \tag{2}$$

where $\{\hat{L}_i^{(k)}\}_{k=1}^K$ denotes the set of $K$ generated candidates for test case $i$. Recovery reflects the model's ability to reconstruct the true linker conditioned on the given fragments.

**Uniqueness** measures the diversity of valid generated samples by computing the fraction of distinct molecules among valid outputs. Let $\mathcal{G}_{valid} \subseteq \mathcal{G}$ denote the subset of valid generated linkers. Then uniqueness is defined as:

$$\text{Uniqueness} = \frac{|\text{Unique}(\mathcal{G}_{\text{valid}})|}{|\mathcal{G}_{\text{valid}}|}, \tag{3}$$

where Unique(·) removes duplicated molecules based on canonical SMILES comparison. This metric evaluates the generative diversity of the model.

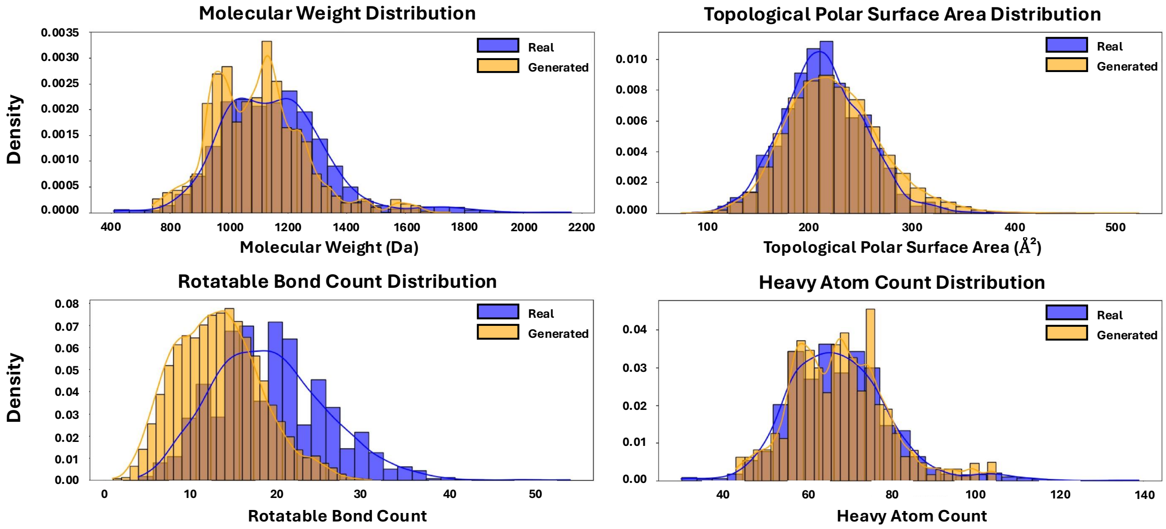


Figure S1: Property distribution gap between generated and true linkers.

### 3 Gated Weights Analysis

In this section we further analyse the relations between the gated weights and different physiochemical properties.

Figure S2 visualizes the evolution of gate weights assigned to different molecular properties along the diffusion trajectory. Several clear temporal patterns can be observed.

First, structure-related attributes such as H-bond donor (HBD) count and rotatable bond (RB) count consistently maintain relatively high weights throughout the denoising process. This suggests that the model persistently relies on these features to constrain geometric flexibility and local interaction patterns, which are critical for maintaining chemically valid linker conformations.

Second, certain global physicochemical properties, such as TPSA and molecular weight, exhibit a gradual increase in gate weight as the time step progresses toward later denoising stages. This trend indicates that, while early steps primarily focus on recovering coarse structural geometry under high noise levels, later stages increasingly emphasize fine-grained physicochemical consistency and global property alignment with the target distribution.

In contrast, properties such as heavy atom count (HA) and H-bond acceptor (HBA) display comparatively moderate and smoother variations over time, implying a more stable but less dominant role during generation.

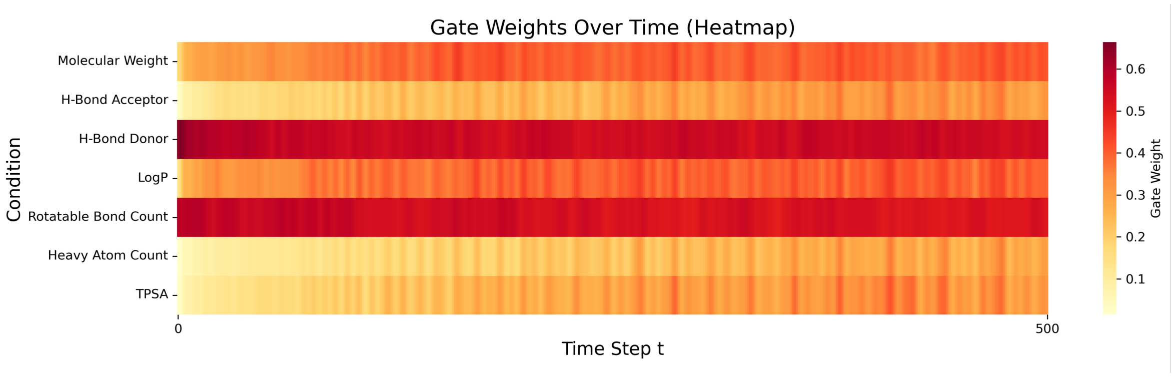


Figure S2: Heatmap visualisation of gate weights overtime

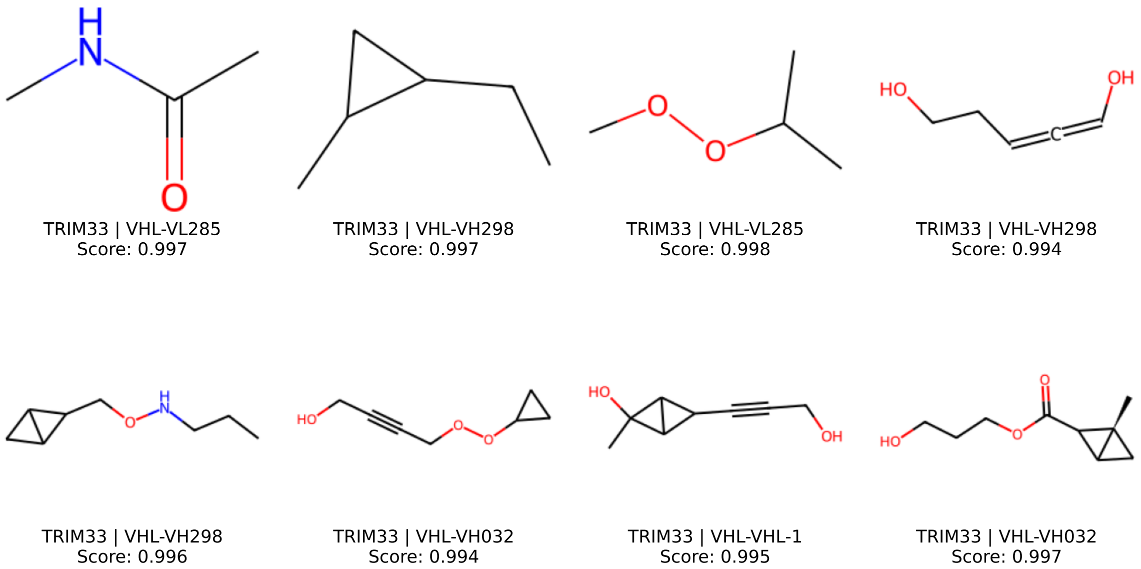


Figure S3: Linker Generation Results of TRIM33.

Overall, the non-uniform and time-dependent weighting patterns confirm that the time-gated fusion mechanism does not act as a static conditioning module. Instead, it dynamically adjusts the contribution of different molecular attributes across diffusion steps, supporting a coarse-to-fine generation process and validating the design of our conditional, time-aware fusion strategy.

### 4 TRIM33-Targeting PROTAC Linker Generation

TRIM33 (Tripartite Motif Containing 33), is a transcriptional co-regulator belonging to the TRIM protein family. It plays a critical role in chromatin remodeling and transcriptional regulation, particularly in TGF-$\beta$ signaling and hematopoietic differentiation. Dysregulation of TRIM33 has been implicated in multiple pathological contexts, including leukemia, tissue fibrosis, and several solid tumors. Despite its biological significance, no PROTACs targeting TRIM33 have been reported to date, making it a compelling candidate for exploratory degrader design.

To evaluate our framework in a realistic design setting, we generated candidate linkers for TRIM33-targeting PROTACs under different VHL-based E3 ligands. The generated candidates were subsequently scored using a pretrained scoring model that estimates structural plausibility and compatibility between the warhead, linker, and E3 ligand. All candidates were ranked according to the predicted score, and the top-scoring molecules are shown in Figure S3.

The top-ranked linkers (scores $\geq 0.994$) display structurally coherent yet chemically diverse architectures consistent with established PROTAC design principles. Most candidates combine flexible aliphatic chains with locally constrained motifs (e.g., small rings or alkynes), achieving a balance between conformational adaptability and geometric stability that is critical for ternary complex formation. The linker lengths are moderate, suggesting that the model implicitly regulates spatial separation between the warhead and the E3 ligand.

Moreover, polar functional groups such as hydroxyl, ether, ester, and amide moieties are frequently observed, contributing to hydrogen-bonding potential and physicochemical compatibility. Despite their diversity, the generated linkers remain chemically reasonable and pharmacologically plausible, indicating that the model effectively captures key structural and property constraints required for TRIM33-targeted PROTAC design.

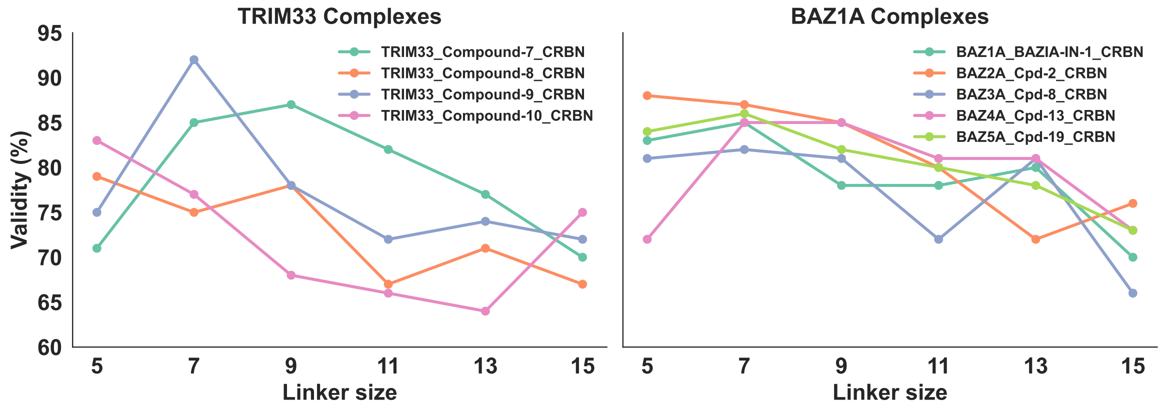


Figure S4: Validity on TRIM33 and BAZ1A with Different Linker sizes.

### 5 Impact of Linker Size on PROTAC Validity

In addition to TRIM33, we further evaluate our model on BAZ1A (Bromodomain Adjacent to Zinc Finger Domain Protein 1A), a chromatin-associated protein involved in transcriptional regulation and DNA damage response. BAZ1A has been implicated in cancer-related pathways and epigenetic regulation, making it an attractive but relatively underexplored target for targeted protein degradation. Including BAZ1A enables us to assess whether the observed trends generalize across distinct protein classes.

Linker length is a critical determinant of PROTAC efficacy. Previous studies have shown that even small variations in linker size can significantly alter ternary complex formation, degradation potency, and selectivity, due to changes in spatial orientation and entropic penalties (e.g., ). An overly short linker may prevent productive protein–protein interaction, whereas an excessively long linker can introduce conformational heterogeneity and reduce complex stability. Therefore, systematic evaluation of linker size is essential for rational PROTAC optimization.

Because our model incorporates molecular property conditioning, and linker size directly influences physicochemical properties (e.g., molecular weight, rotatable bonds, TPSA), we trained a lightweight regression model on the training set to estimate and adjust the property shift induced by varying linker sizes. This adjustment ensures that differences in validity across sizes are not confounded by systematic property deviations, enabling a more rigorous evaluation of linker length effects.

Figure S4 summarize the validity (%) of generated PROTACs across linker sizes (5–15) for TRIM33 and BAZ1A complexes. Several consistent trends emerge.

For TRIM33 complexes, validity generally peaks at intermediate linker sizes (7-9). For example, TRIM33_Compound-7_CRBN achieves its highest validity at size 9 (87%), while TRIM33_Compound-9_CRBN peaks at size 7 (92%). In contrast, larger linkers (13–15) tend to reduce validity across most complexes. This pattern aligns with the notion that moderate linker lengths better balance spatial reach and conformational stability.

For BAZ1A complexes, a similar trend is observed. Most compounds exhibit highest validity at sizes 7–9 (e.g., BAZ2A_Cpd-2_CRBN reaches 87% at size 7, BAZ4A_Cpd-13_CRBN maintains 85% at sizes 7 and 9). Larger sizes ($\geq$13) generally show decreased validity, particularly for BAZ3A_Cpd-8_CRBN and BAZ1A_BAZIA-IN-1_CRBN. Notably, the performance curves across targets display comparable shapes, suggesting that the effect of linker size follows a consistent structural principle rather than being target-specific.

Overall, the results indicate that intermediate linker lengths (7–9) yield the most stable and valid generations across both targets. Extremely short or long linkers reduce generation validity, likely due to geometric incompatibility or excessive flexibility. These findings are consistent with established PROTAC design knowledge and further demonstrate that our framework can sensitively capture linker-size-dependent structural constraints in a target-agnostic manner.

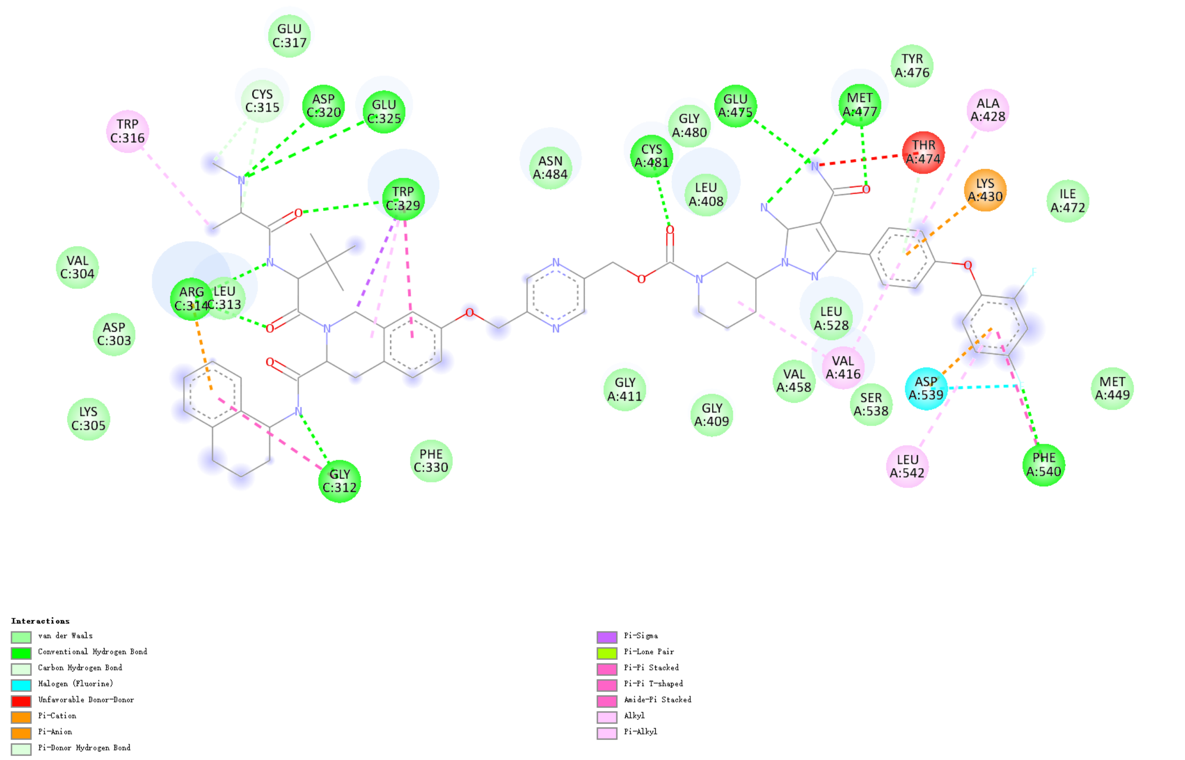


Figure S5: Protein–PROTAC interaction map of the 6W7O complex, illustrating hydrogen bonds, van der Waals, $\pi$ interactions, hydrophobic contacts and so on between the BCPyr PROTAC and the surrounding residues in the binding interface.

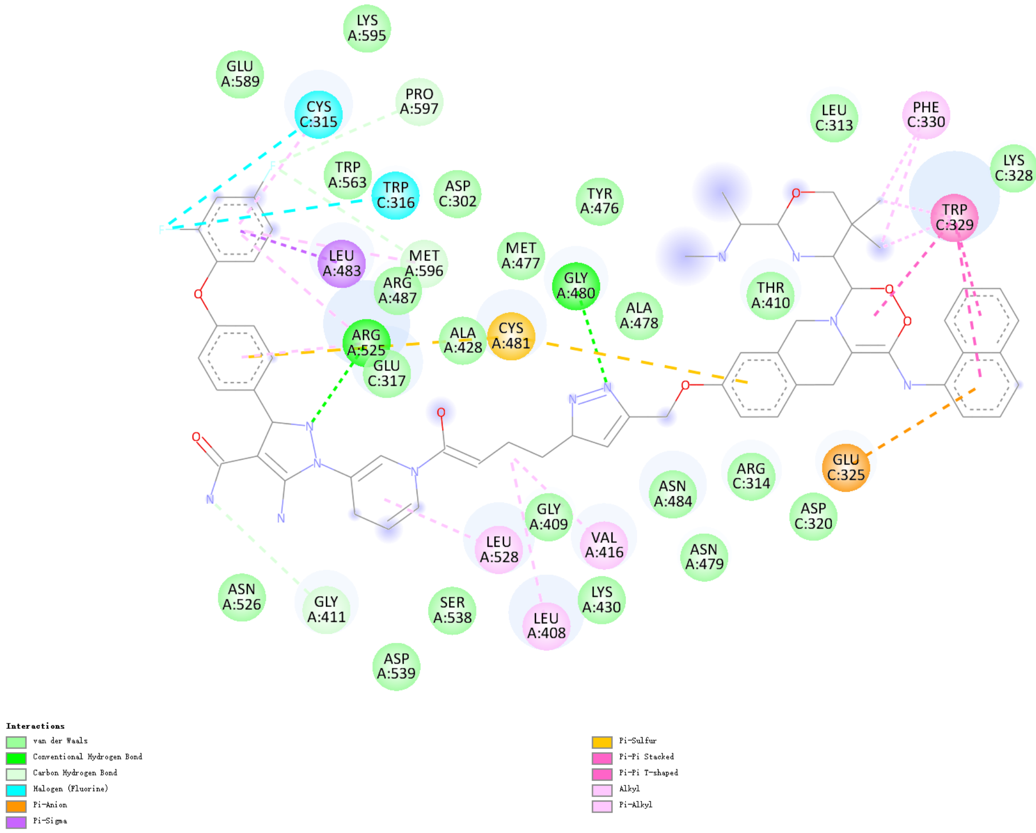


Figure S6: Protein–PROTAC interaction map of the generated complex, illustrating hydrogen bonds, van der Waals, $\pi$ interactions, hydrophobic contacts and so on between the DiffPROTAC generated PROTAC and the surrounding residues of 6W7O in the binding interface.

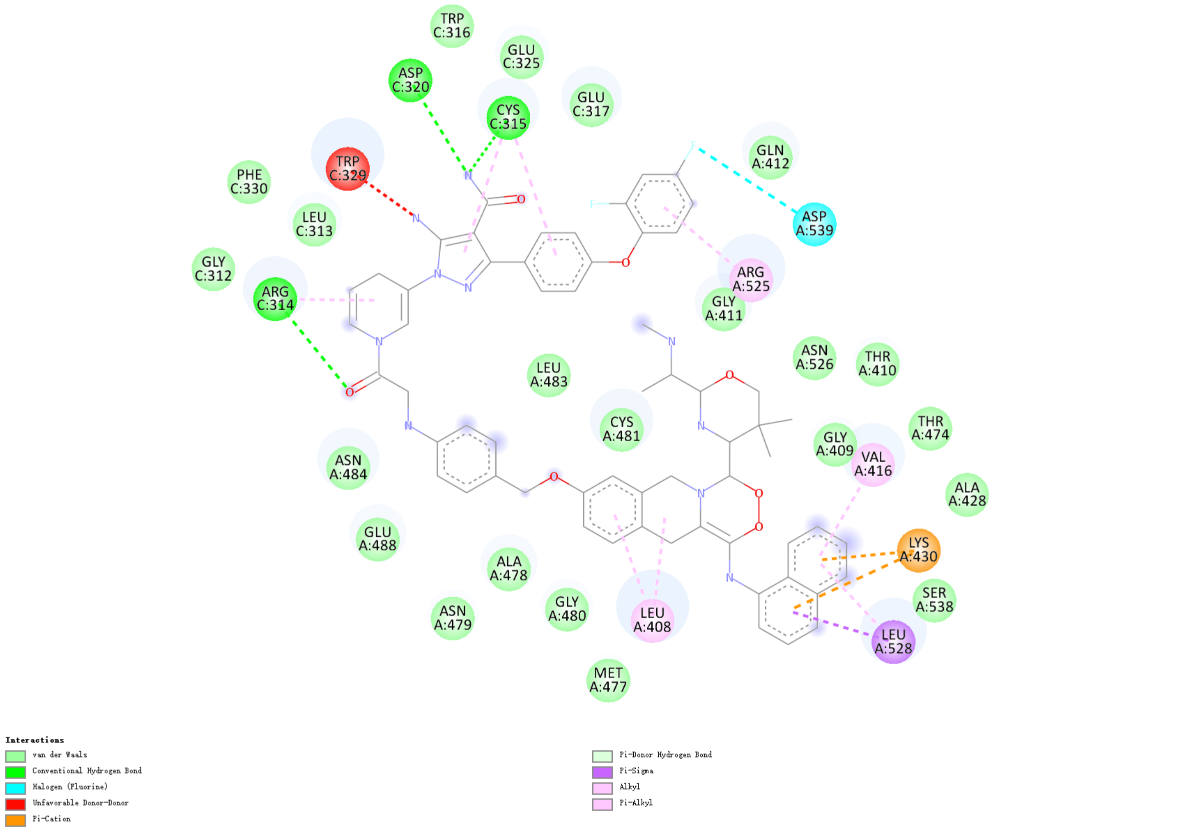


Figure S7: Protein–PROTAC interaction map of the generated complex, illustrating hydrogen bonds, van der Waals, $\pi$ interactions, hydrophobic contacts and so on between the DesignMaster generated PROTAC and the surrounding residues of 6W7O the binding interface.
